# Evaluation of AlphaFold3 for the fatty acids docking to human fatty acid-binding proteins

**DOI:** 10.1101/2024.07.25.605175

**Authors:** Ki Hyun Nam

**Author notes:** Corresponding author (K.H. Nam).

## Abstract

Human fatty acid-binding proteins (FABPs) are involved in many aspects of lipid metabolism, such as the uptake, transport, and storage of lipophilic molecules, as well as cellular functions. Understanding how FABPs recognize fatty acids (FAs) is crucial for identifying FABP function and applications, such as in inhibitor design or biomarker development. The recently developed AlphaFold3 (AF3) demonstrates significantly higher accuracy than other prediction tools, particularly in predicting protein–ligand interactions with state-of-the-art docking tools. Studies on whether AF3 can be used to identify the FAs of FABP are lacking. To assess the accuracy of FA docking to FABPs using AF3, models of FA docked into FABP generated using AF3 were compared with experimentally determined FA-bound FABP structures. FA ligands in AF3 structures docked reliably into the FA-binding pocket of FABPs; however, the detailed binding configuration of most FA ligands docked into FABPs and the interaction between FA and FABP determined using AF3 and experimentally differed. These results will aid in understanding FA docking to FABPs and other FA-binding proteins using AF3.

## 1. Introduction

Fatty acids (FAs)—an important energy resource—are involved in complex cellular functions, such as metabolic regulation and responses, enzymatic and transcriptional gene expression regulation, cell growth and survival pathways, and inflammatory responses [1–3]. FA-binding proteins (FABPs) are intracellular lipid chaperones that coordinate lipid responses in cells and are strongly linked to metabolic, inflammatory, and energy homeostasis pathways [4–8].

In humans, 10 FABPs (FABP1–FABP9 and FABP12) have been identified, exhibiting specific expression patterns in various tissues, such as the liver, intestine, heart, adipocyte, epidermis, ileum, brain, myelin, and testis [9]. FABPs are involved in various lipid metabolism processes, such as the uptake, transport, storage of lipophilic molecules (e.g., long-chain saturated FAs, unsaturated FAs, and other lipids) [10]. Human FABP encodes a ∼15 kDa protein, and the amino acid sequence identities among 10 human FABPs vary in the range of 15%–70% [11]. Despite the wide variance in amino acid sequences, FABPs commonly exhibit a β-barrel structure [12]. Hydrophobic ligands, such as saturated or unsaturated long-chain FAs, lipids, or lipophilic molecules, noncovalently and reversibly bind to the inside of the β-barrel folds of FABPs [13, 14]. Each FABP exhibits significant differences in ligand selectivity, binding affinity, and binding mechanism [15]. FABPs bind not only to lipophilic ligands to perform unique functions within specific tissues but also to other FAs. For example, FABP7, expressed in the brain, preferentially binds docosahexaenoic acid but can also bind other FAs, such as oleic acid (OLA) and palmitic acid (PLM) [16, 17]. Understanding FA recognition by FABPs will not only help to understand their functions but also provide structural information that may contribute to the development of inhibitors or biomarkers for specific FABPs [18, 19].

Protein structure prediction by AlphaFold2 has revolutionized protein structure and interaction modeling and is widely applied in protein modeling and design [20]. Recently, AlphaFold3 (AF3) was introduced with a significantly updated diffusion-based architecture that can predict the binding structures of complexes containing proteins, nucleic acids, small molecules, and ions [21]. In particular, AF3 provides significantly higher accuracy in predicting protein–ligand interactions than state-of-the-art docking tools [21].

Several crystal structures of FA-bound human FABPs (FABP1–9) have been determined, but the FA recognition mechanisms have not been fully elucidated. AF3 can be used to study FA substrate docking on FABPs and could yield valuable insights into how FABPs recognize FAs. However, the reliability of AF3 in providing accurate information has not been fully verified.

In this study, the structures of FA docked on FABPs were generated using AF3. AF3 predicted models of FA binding on FABPs were analyzed and compared with experimental structures of FABPs complexed with FAs. Structural comparisons between AF3 predictions and experimental data showed the utility and limitations of using AF3 for FA docking on FABPs. This study provides valuable insights into FABP substrate binding and offers guidance on how to effectively utilize AF3.

## 2. Material and methods

### 2.1. Selection and preparation of FABPs

Canonical amino acid sequences of FABP1–5 and FABP7–9 were obtained from UniProt (http://uniprot.org) [22] for the AF3 docking study. The experimentally determined structures of FABP1–5 and FABP7–9 complexed with FAs (myristic acid [MYR], palmitic acid [PLM], or oleic acid [OLA]) were obtained from Protein Data Bank (http://rcsb.org) [23].

### 2.2. Molecular docking

FAs (MYR, PLM, or OLA) were docked into FABPs in the AF3 server (https://golgi.sandbox.google.com/) [21]. For FABP1, FAs were automatically docked into FABP1 with one or two binding modes. For other FABPs, one FA molecule was automatically docked into FABPs. AF3 provided five docking models for FA docking into FABP, and these structures were used for structural analysis. The model structures of FA-docked FABP using AF3 have been deposited in Zenodo (https://doi.org/10.5281/zenodo.12747397).

### 2.3. Structure comparison and bioinformatics

Atom-level structural comparison of FA molecules from AF3 and experimentally determined structures were aligned with PyMOL (http://pymol.org). The FA–FAPB interaction was analyzed using the AF3 model with the highest interface predicted template modeling (ipTM) scores and the experimental structure with the highest resolution. Interactions between FABPs and ligands were analyzed using Protein–Ligand Interaction Profiler [24]. Binding affinity between FABP and FA were calculated with the PRODIGY-LIGAND web server [25]. The structure were visualized with PyMOL.

## 3. Results

### 3.1. Data preparation

Three FAs: myristic acid (MYR), PLM, and OLA, are available for docking on the AF3 server. The docking of FA ligands into FABP using AF3 (AF3-FABP) was compared with experimentally determined structures of FABPs (EXP-FABP), such as MYR-, PLM-, or OLA-bound FABP1–5 and FABP7–9. Structural comparisons between AF3 and experimental structures for FABP6 and FABP12 were excluded because their experimental structures complexed with FA have not been determined. In some FABP structures, specific amino acids were substituted or small molecules were bound to the FA-binding pocket of FABPs, affecting the binding and conformation of FA [26]. For example, in PLM–FABP1 (PDB codes: 7DZE, 7DZF, 7DZG, 7DZH, 7DZI, 7DZJ, 7DZK, and 7DZL), the amino acid Phe63 of FABP1, located in the FA-binding pocket, was substituted with a xanthone amino acid for time-resolved serial crystallography [26], affecting the binding configuration of PLM. In the MYR– FABP4 mutant (PDB codes: 7FYH and 7G0X), 6-fluoro-1,3-benzothiazol-2-amine or isoquinolin-3-amine were bound to the FA-binding site, affecting the binding configuration of MYR (unpublished). Thus, experimental FA–FABP structures that affected the FA-binding pocket were excluded from this study (see Discussion).

FA ligand docking into FABP using AF3 automatically generated five protein–ligand docking models. AF3 provides confidence metrics, including the predicted template modeling (pTM) and the interface predicted template modeling (ipTM) scores [21]. A pTM score of >0.5 indicates that the overall predicted fold for the complex might be similar to the true structure. The ipTM score measures the accuracy of the predicted relative positions of the subunits within the complex. An ipTM value of >0.8, 0.6–0.8, and <0.6 represents confident high-quality predictions, correct or incorrect predictions, and failed predictions, respectively. Each highest scored FABP–FA docking structure generated using AF3 showed an ipTM score of >0.86 and a pTM score of >0.89, indicating that the docking results are reliable, based on the standard criteria of AF3 (Figure 1). The calculated binding affinities between FABP and FA were similar between the five AF3-FABP model structures and EXP-FABP, with highest resolution structure.

**Figure 1.**
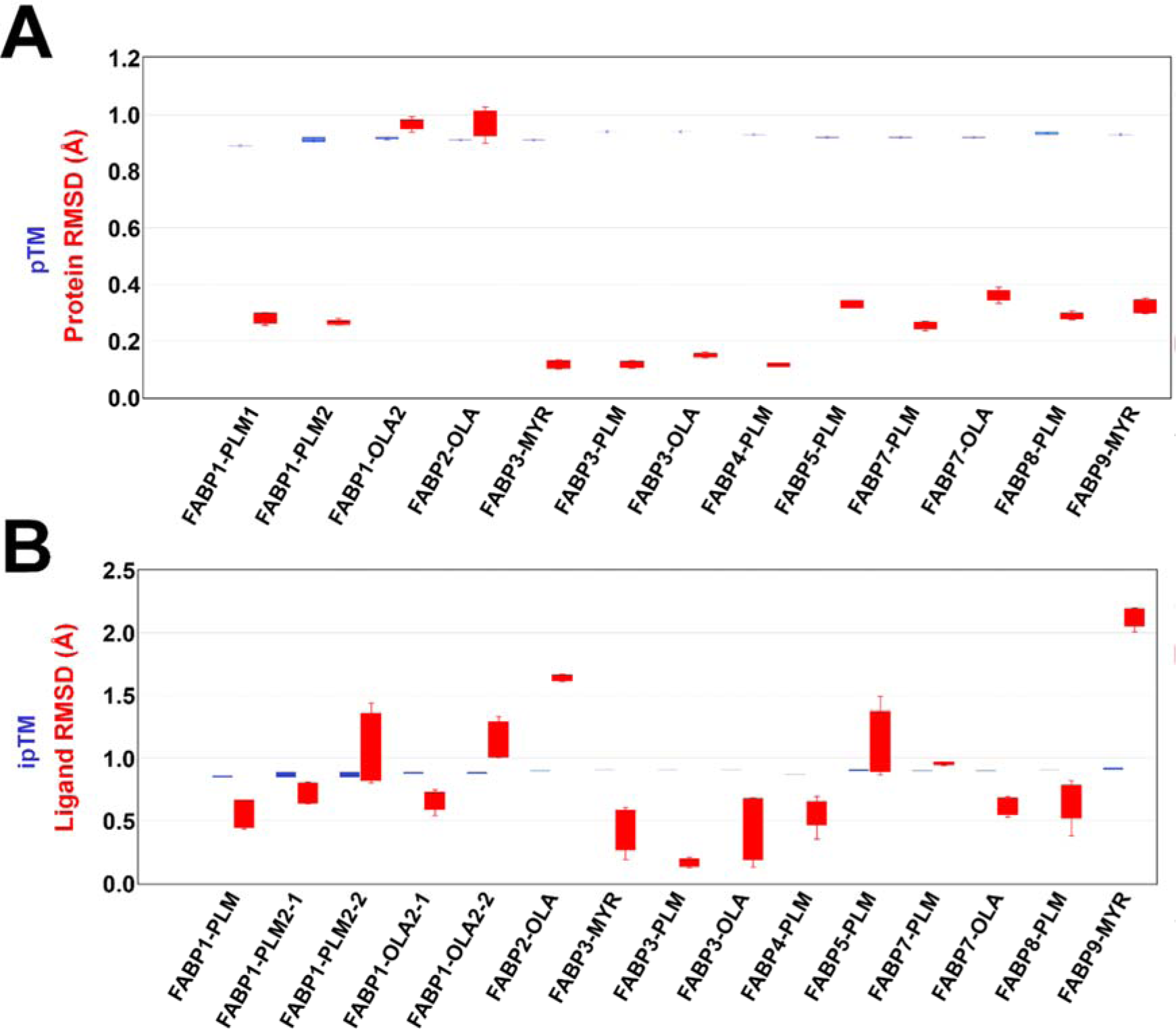
Protein and ligand RMSD values between AF3 and experimental FABP structures. (A) Comparison of AF3 and experimental FABP structures. The five pTM scores generated using AF3 and RMSD values between AF3 and experimental FABP structures are indicated by blue and red bars, respectively. (B) Comparison of AF3 and experimental FA structures. The five ipTM scores generated using AF3 and the RMSD values between AF3 and experimental FA structures are indicated by blue and red bars, respectively. The FABP and FA structures generated using AF3 were compared with experimental structures of FABP1–PLM (PDB code: 3STM), FABP1– PLM2 (3VG7), FABP1–PLA2 (2LKK), FABP2–OLA (2M05), FABP3–MYR (4TKH), FABP3–PLM (7FFK), FABP3– OLA (7WE5), FABP4–PLM (2HNX), FABP5–PLM (1B56), FABP7–PLM (7E25), FABP7–OLA (1FE3), FABP8– PLM (6S2M), and FABP9–MYR (7FY1).

To evaluate the FABP and FA structures generated using AF3, root mean square deviation (RMSD) values between AF3-FABP and the EXP-FABP structure with highest resolution were calculated. All AF3-FABP structures showed structural similarity to EXP-FABPs with RMSD values of <0.4 Å, excluding the two FA-bound structures of EXP-FABP1–PLM2 and EXP-FABP2–OLA that had RMSD values of >0.9 Å (Figure 1A). The five FA ligands docked into FABP with AF3 were compared with experimentally determined FA in FABP with highest resolution structures. FA structures from FABP1–PLM, FABP3–MYR, FABP3–PLM, FABP3– OLA, FABP4–PLM, FABP7–PLM, FABP7–OLA, and FABP8–PLM generated using AF3 had RMSD values of <0.9 Å compared with the experimental FA structures. By contrast, FA structures from FABP1–PLM2, FABP2–OLA, FABP5–PLM, and FABP9–MYR generated using AF3 had RMSD values of >0.9 Å compared with the experimental FA structures. Taken together, the pTM and ipTM scores generated using AF3 did not correlate with experimental data; some FABP–FA structures generated with AF3 or experimentally differed significantly (see below).

### 3.2. Structural comparison of FA docked into FABP

#### 3.2.1. FABP1

FABP1 (liver-type FABP or L-FABP) is expressed mainly in the liver but is also expressed in the intestine and kidney, participating in FA metabolism in the cytoplasm [13, 27]. FABP1 binds various lipophilic molecules, such as long-chain FAs, fatty acyl-CoAs, peroxisome proliferators, prostaglandins, bile acids, bilirubin, and hydroxy and hydrophobic ligands [12, 28]. FABP1 is unique from other FABPs in that two FA ligands can bind to the FA-binding site [29]. In PDB, the crystal structures of one or two PLM–bound FABP1 and two OLA-bound FABP1 have been deposited, which were compared with the FA-docked AF3-FABP1 structure.

In one PLM-docked AF3-FABP1, all PLM models with a U-shaped conformation were docked into the FA-binding site of FABP1 (Figure 2A). The middle of the aliphatic chain of PLM was located inside the FA-binding pocket, and the tail of the aliphatic chain and the carboxyl group of PLM were toward the substrate-binding entrance of FABP1. The position and conformation of PLM in AF3-FABP1 is similar with those of PLM in EXP-FABP1 (PDB code: 3STM) (Figure 2B and 2C). The superimposition of PLM from AF3-FABP1 and EXP-FABP1-3STM has an RMSD of 0.427–0.661 Å. In AF3-FABP1 with one PLM, the carboxyl group of PLM1 interacted with Ser39 (3.06 Å), Asn111 (3.54 Å), Arg122 (2.77 and 3.26 Å), and Ser124 (3.03 Å), and the aliphatic chain of PLM1 was stabilized by hydrophobic interactions with Phe50 (3.89 and 3.93 Å) and Leu71 (3.99 Å) (Figure 2D). In EXP-FABP1 with one PLM, the carboxyl group of PLM1 interacted with Ser39 (2.72 Å) and Arg122 (2.97 and 3.27 Å) and stabilized through water bridges with Ser39 (3.77 Å) and Asn111 (3.72 Å). The aliphatic chain of PLM1 was stabilized by hydrophobic interactions with Phe50 (3.62 Å), Phe63 (3.94 Å), Leu91 (3.93 Å), and Thr102 (3.50 Å) (Figure 2E). The superimposition of one PLM-bound AF3-FABP1 and EXP-FABP1 showed that the conformation of the PLM ligands and some of PLM-interacting amino acids were different (Figure 2F), but the overall structure had high similarity with an RMSD of 0.255–0.293 Å.

**Figure 2.**
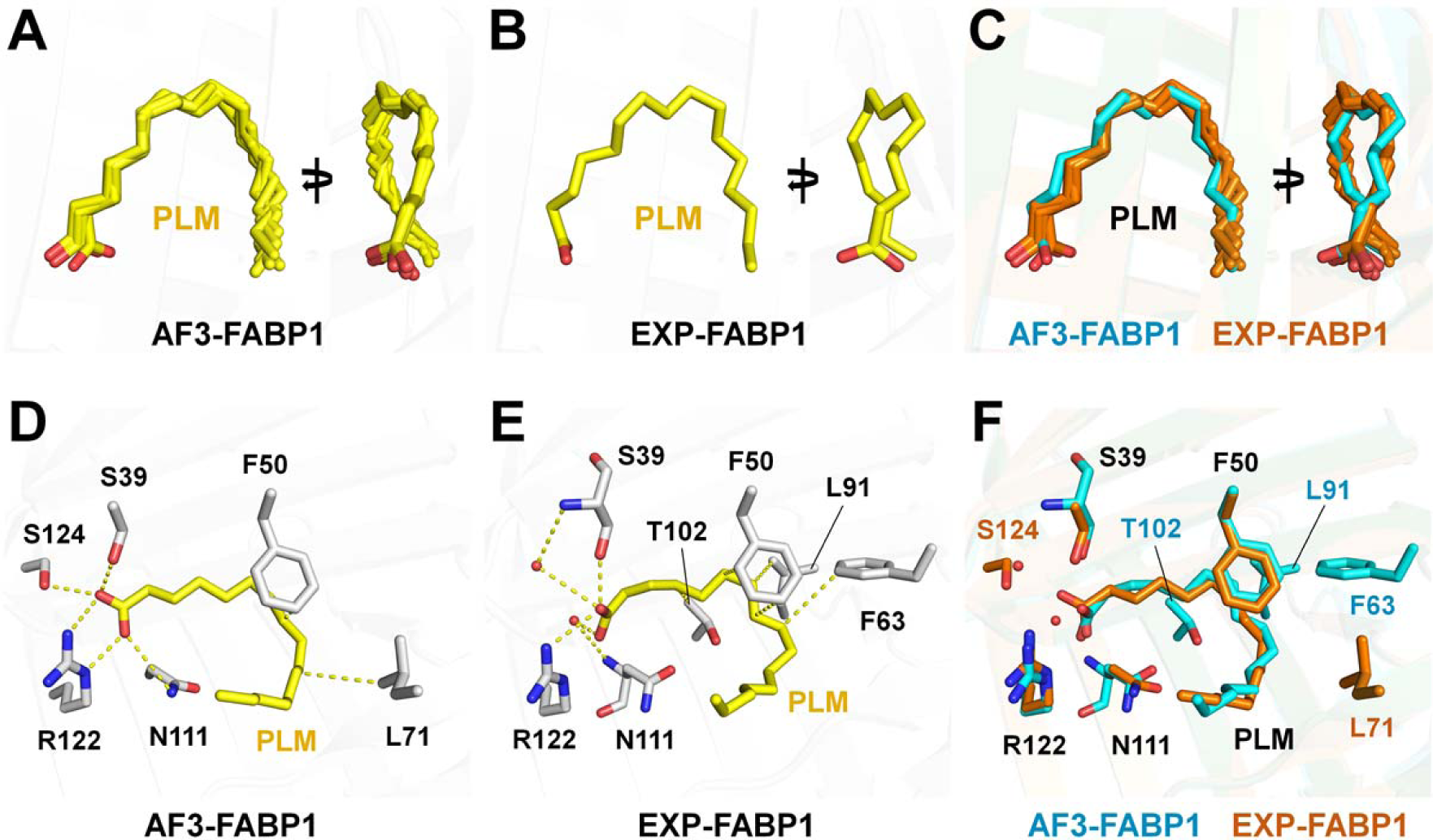
Structural comparison of one PLM in AF3-FABP1 and EXP-FABP1. (A) Superimposition of five PLM ligands from AF3-FABP1. (B) PLM in the EXP-FABP1 structure (PDB code: 3STM). (C) Superimposition of PLM ligands from the AF3-FABP1 (orange) and EXP-FABP1 (cyan) structures. Interactions of PLM with the (D) AF3-FABP1 and (E) EXP-FABP1 (3STM) structures. (F) Superimposition of the PLM-binding site of AF3-FABP1 and EXP-FABP1.

In the two PLM-docked AF3-FABP1 structure, the first PLM (PLM1) with a U-shaped conformation was docked inside the FA-binding pocket of FABP1 (Figure 3A). This binding configuration was mostly similar with that of FABP1 with one PLM-binding mode (Figure 2A). The second PLM (PLM2) with a relatively linear conformation was positioned near the PLM1 and the entrance of the FA-binding pocket (Figure 3A). Notably, among the five PLM2 models generated using AF3, the carboxyl groups of the two PLM2 models of FABP1 were oriented inside the FA-binding pocket, whereas the carboxyl groups of the other three PLM2 models were oriented toward the entrance of FA-binding pocket (Figure 3A). In 12 two PLM-bound EXP-FABP1 structures (PDB codes: 3B2H, 3B2I, 3B2J, 3B2K, 3B2L, 3STK, 3VG2, 3VG3, 3VG4, 3VG5, 3VG6, and 3VG7), the position and conformation of PLM1 was similar with PLM1 of AF3-FABP1. All carboxyl groups of PLM2 of AF3-FABP1 were oriented toward PLM1 located on the inside of the FA-binding pocket (Figure 3B). These results indicate that the three PLM2, oriented with the carboxyl group toward the entrance of the FA-binding pocket, docked using AF3-FABP1, differed from the experimental results. The superimposition of PLM1 and PLM2 from AF3-FABP1 and EXP-FABP1 (3STM) showed RMSD values of 0.632–0.808 and 0.799–1.438 Å, respectively. The position of PLM1 and PLM2 from AF3-FABP1 and EXP-FABP1 showed similarity. Considering that various conformations of PLM1 and PLM2 exist among the 12 EXP-FABP1 structures, the conformations of two PLM ligands docked in FABP1 using AF3 are considered reasonable docking (Figure 3C). In the two PLM-bound FABP1 structures, the orientation of the top three scored PLM2 models in AF3-FABP1 differed from that of PLM2 on EXP-FABP1. Therefore, the protein–ligand interactions of AF3-FABP1 were analyzed using the fourth highest scored AF3 model, whose ligand orientation is identical to the experimental structure. In the two PLM docked AF3-FABP1 structure, the carboxyl group of PLM1 interacted with Ser39 (2.59 Å), Arg122 (2.86 and 2.91 Å), and Ser124 (3.46 Å), and the aliphatic chain of PLM1 was stabilized by hydrophobic interactions with Ile41 (3.96 Å), Phe50 (3.79 Å), and Leu71 (3.85 Å) (Figure 3D).The carboxyl group of PLM2 interacted with Asn111 (2.79 Å) and Arg122 (3.16 Å), and the aliphatic chain of PLM2 was stabilized by hydrophobic interactions with Leu24 (3.74 Å) and Ile52 (3.52 Å) (Figure 3D). In the two PLM-bound EXP-FABP1 structure (PDB code: 3VG7), the carboxyl group of PLM1 interacted with Ser39 (2.60 Å), Arg122 (2.82 and 2.93 Å), and Ser124 (3.50 Å) and was stabilized by water bridges with Asn111 (3.99 Å) and Arg122 (3.40 Å). The aliphatic chain of PLM1 was stabilized by hydrophobic interactions with Phe50 (3.74 Å), Phe63 (3.98 Å), Leu91 (3.99 Å), Phe95 (3.40 Å), Thr102 (3.70 Å), Ile109 (3.92 Å), and Met113 (3.91 Å) (Figure 3E). The carboxyl group of PLM2 interacted with Arg122 (3.28 Å) and the water bridge with Asn111 (3.68 Å). The aliphatic chain of PLM2 was stabilized by hydrophobic interactions with Leu24 (3.67 Å), Lys31 (3.76 Å), Ile35 (3.90 Å), and Ile52 (3.87 Å) (Figure 3E). The superimposition of the two PLM-bound AF3-FABP1 and EXP-FABP1 structures revealed that the positions of the PLM ligands were similar. However, in EXP-FABP1, the two PLM molecules had more extended interactions with FABP1 than the PLM in AF3-FABP1 (Figure 3F).

**Figure 3.**
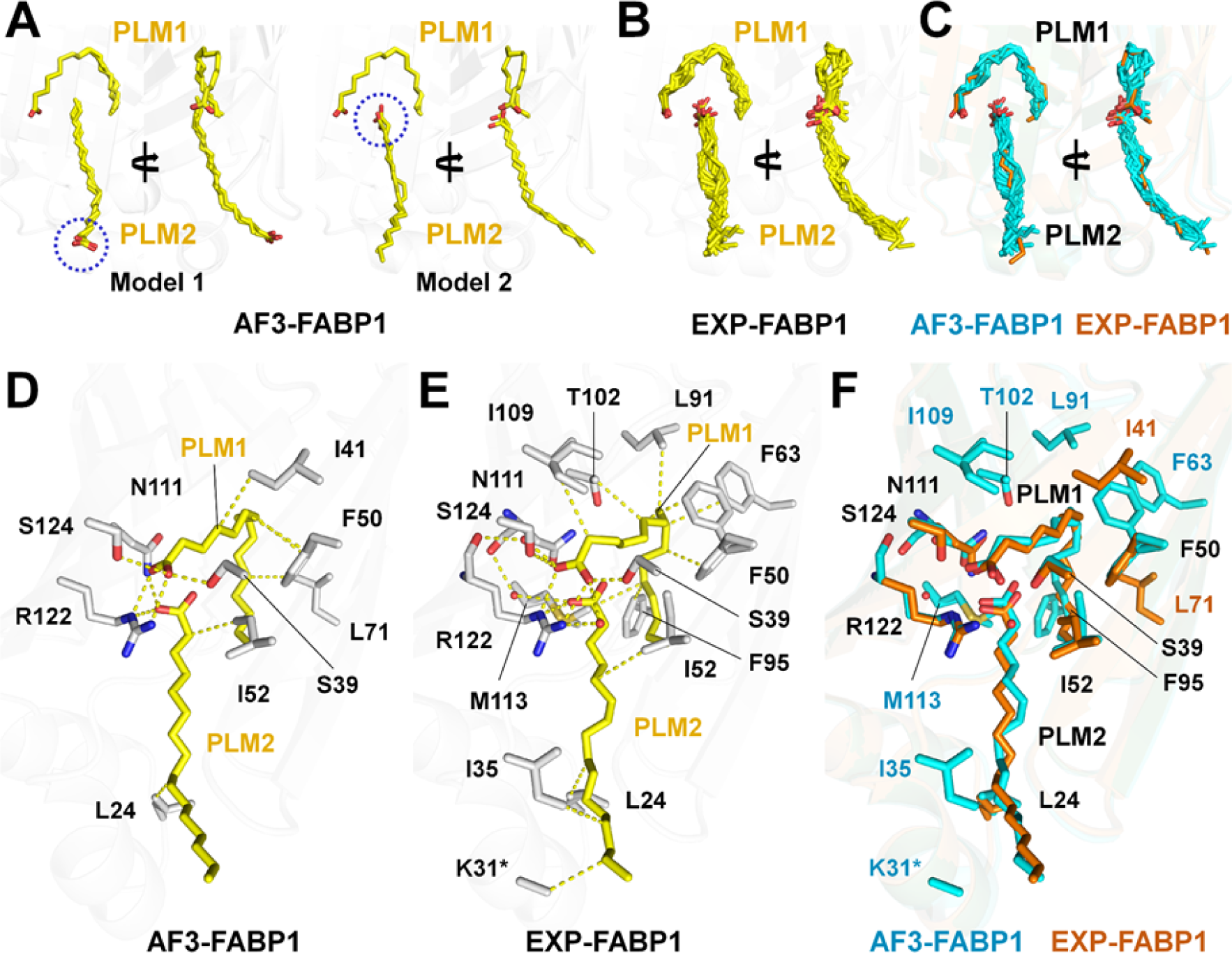
Structural comparison of two PLM in AF3-FABP1 and EXP-FABP1 structures. (A) Superimposition of five PLM1 and PLM2 ligands from AF3-FABP1. (B) PLM on the EXP-FABP1 structure (PDB code: 3B2H, 3B2I, 3B2J, 3B2K, 3B2L, 3STK3VG2, 3VG3, 3VG4, 3VG5, 3VG6, and 3VG7). (C) Superimposition of PLM ligands from AF3-FABP1 (orange) and EXP-FABP1 (cyan) structures. Interactions of PLM ligands with (D) AF3-FABP1 and (E) EXP-FABP1 (PDB code: 3VG7) structure. (F) Superimposition of the two PLM-binding sites of AF3-FABP1 and EXP-FABP1. Some coordinates for the side chain of K31* of EXP-FABP1 were missing in PDB.

In the two OLA2-bound AF3-FABP1 structure, the first OLA (OLA1) molecule with a U-shaped conformation was located inside of FA binding, and the second OLA (OLA2) with a relatively linear conformation was located from U-shaped OLA1 and entrance of FA-binding pocket (Figure 4A). Notably, in the five OLA2 models, the carboxyl groups of the four OLA2 were oriented toward the FA-binding entrance, whereas the carboxyl groups of other one OLA2 were oriented toward the OLA1 located in the FA-binding pocket (Figure 4A). In the two OLA-bound EXP-FABP1 structures (PDB code: 2LKK), the carboxyl groups of OLA2 were oriented toward the FA-binding entrance (Figure 4B). Superimposition of two PLM-bound AF3-FABP1 and EXP-FABP1 showed the distinct position of PLM1 and PLM2 (Figure 4C). PLM1 and PLM2 from AF3-FABP1 and EXP-FABP1 had RMSD values of 0.537–0.744 and 1.004–1.333 Å, respectively. In the two OLA-docked AF3-FABP1, the carboxyl group of OLA1 interacted with Ser39 (2.49 Å), Arg122 (2.88 and 2.89 Å), and Ser124 (3.64 Å), and the aliphatic chain of OLA1 was stabilized by hydrophobic interactions with Phe50 (3.78 Å), Ile59 (3.85 Å), Phe63 (3.84 Å), Leu71 (3.92 Å), and Phe95 (3.77 Å) (Figure 4D). The carboxyl group of OLA2 interacted with Ser56 (3.05 Å), and the aliphatic chain of OLA2 was stabilized by hydrophobic interactions with Phe15 (3.64 Å), Leu24 (3.92 Å), Ala54 (3.90 Å), and Lys57 (3.95 Å) (Figure 4D). In the two OLA-bound EXP-FABP1 structure (PDB code: 2LKK), the carboxyl group of OLA1 interacted with Ser39 (2.38 Å), Arg122 (2.77 Å), and Ser124 (2.37 Å), and the aliphatic chain of OLA1 was stabilized by hydrophobic interactions with Ile41 (3.81 Å), Ile52 (3.86 Å), Leu71 (3.68 Å), Phe95 (3.65 Å), Thr102 (3.93 Å), and Ile109 (3.93 Å) (Figure 4E). The carboxyl group of OLA2 interacted with Lys31 (3.40 Å) and Ser56 (2.37 Å), and the aliphatic chain of OLA2 was stabilized by hydrophobic interactions with Phe15 (3.48 and 3.94 Å), Leu24 (3.81 Å), Leu28 (3.65 Å), Ile52 (3.92 Å), and Phe120 (3.89 Å) (Figure 4E). The superimposition of the two OLA-bound AF3-FABP1 and EXP-FABP1 structures exhibited the distinct interaction between the aliphatic chain of PLM2 and FABP1 (Figure 4F).

**Figure 4.**
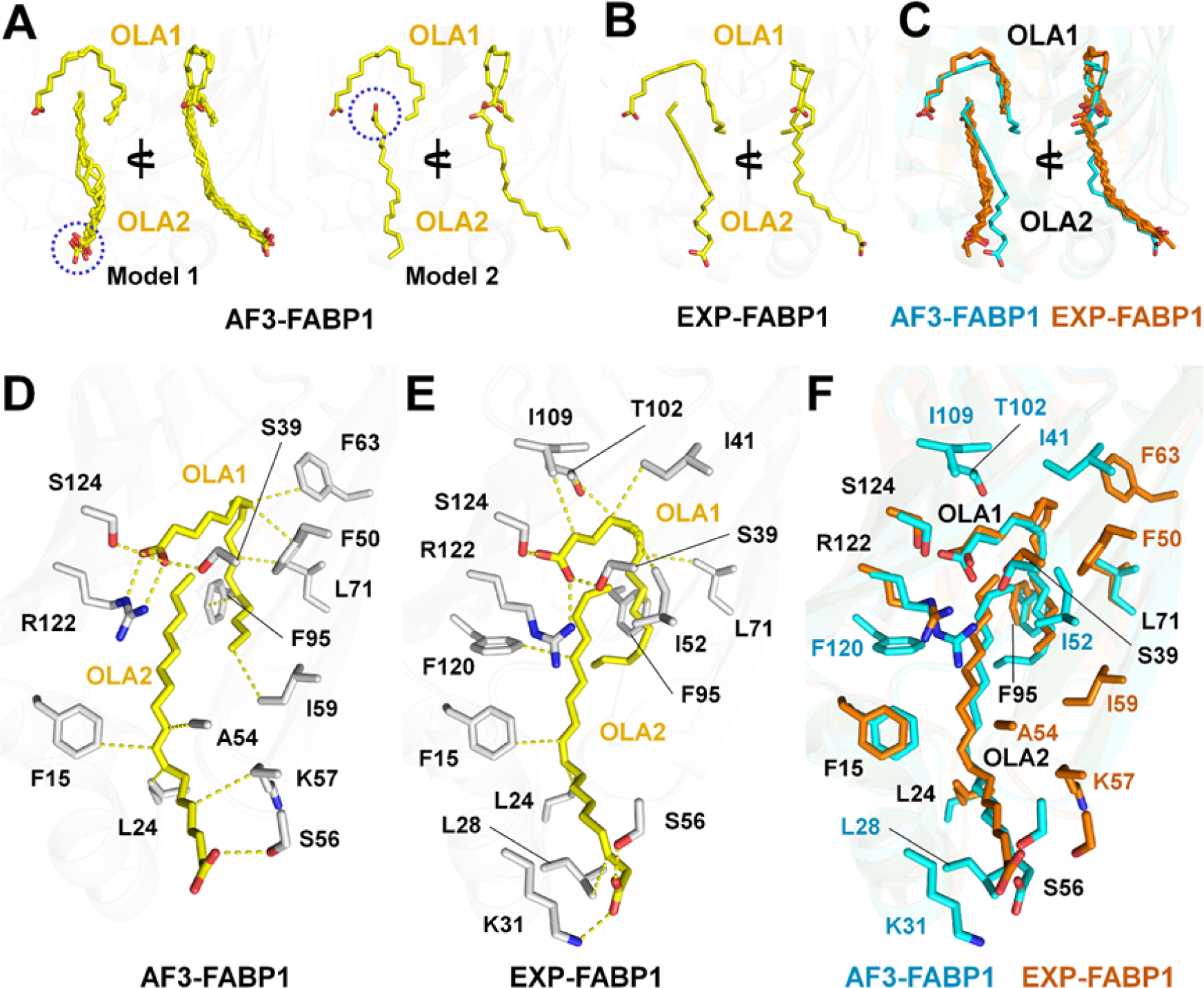
Structural comparison of the two OLA in AF3-FABP1 and EXP-FABP1 structures. (A) Superimposition of the five OLA1 and OLA2 ligands from AF3-FABP1. (B) OLA on EXP-FABP1 structure (PDB code: 2LKK). (C) Superimposition of OLA ligands from the AF3-FABP1 (orange) and EXP-FABP1 (cyan) structures. Interactions of OLA ligands with the (D) AF3-FABP1 and (E) EXP-FABP1 (PDB code: 2LKK) structures. (F) Superimposition of the two OLA-binding sites of AF3-FABP1 and EXP-FABP1.

#### 3.2.2. FABP2

FABP2 (intestinal FABP or I-FABP) is primarily expressed in the small intestine as well as in the liver [30]. FABP2 is involved in dietary lipid absorption and lipid uptake [31]. In PDB, the crystal structure of FABP2–OLA (PDB code: 2M05) has been deposited. The structures of OLA-docked FABP2 generated using AF3 and experimentally determined OLA-bound FABP2 were compared.

In AF3-FABP2 and EXP-FABP2, OLA was commonly located on the FA-binding site of FABP2. The carboxyl group of OLA was toward the inside of the FA-binding pocket, and the tail of the aliphatic chain of OLA was toward the FA-binding entrance (Figure 2A and 2B). Notably, the binding position and configuration of OLA between AF3 and experimental structures were significantly different. In AF3-FABP2, OLA exhibited a C-shaped conformation with a *cis*-conformation at the double bond between C9 and C10 (Figure 5A). The four OLA models exhibited bond angles of 117°–118.6° and 118.0°–120.4° at C8–C9– C10 and C9–C10–C11, respectively, whereas the other one OLA model exhibited bond angles of 172.3° and 173.4° at C8–C9–C10 and C9–C10–C11, respectively. In EXP-FABP2, OLA exhibited several curved-shaped conformations with a *trans*-configuration at the double bond at C9–C10 (Figure 5B). This OLA structure exhibited bond angles of 120.2° and 120.1° at C8–C9–C10 and C9–C10–C11, respectively. The superimposition of OLA from AF3-FABP2 and EXP-FABP2 had an RMDS value of 1.609–1.671 Å, showing the significantly different position of OLA on FABP2 (Figure 5C). The distances between the carboxyl group, C19 atom, and the tail of the aliphatic chain of OLA in AF3-FABP2 and EXP-FABP2 were 2.7, 6.9, and 3.5 Å, respectively (Figure 5C). In AF3-FABP2, the carboxyl group of OLA interacted with Arg107 (2.69 and 2.77 Å), and the aliphatic chain of OLA was stabilized by hydrophobic interactions with Phe18 (3.78 Å), Lys28 (4.00 Å), Phe56 (3.76 Å), Tyr71 (3.76 Å), Leu73 (3.87 Å), Ala74 (3.92 Å), Leu79 (3.93 Å), Phe94 (3.59 Å), Leu103 (3.88 Å), and Tyr118 (3.36 Å) (Figure 5D). In EXP-FABP2 (PDB code: 2MO5), the carboxyl group of OLA interacted with Trp83 (3.96 Å), Arg107 (2.82 and 2.88 Å), and Gln116 (3.68 Å), and the aliphatic chain of OLA was stabilized by hydrophobic interactions with Phe56 (3.25 Å), Tyr71 (3.40 Å), and Phe94 (3.45 Å) (Figure 5E). The superimposition of the OLA-binding site of AF3-FABP2 and EXP-FABP2 showed that the residues involved in forming the FA-binding pocket and in FA interactions differed significantly due to the different position and conformation of OLA (Figure 5F).

**Figure 5.**
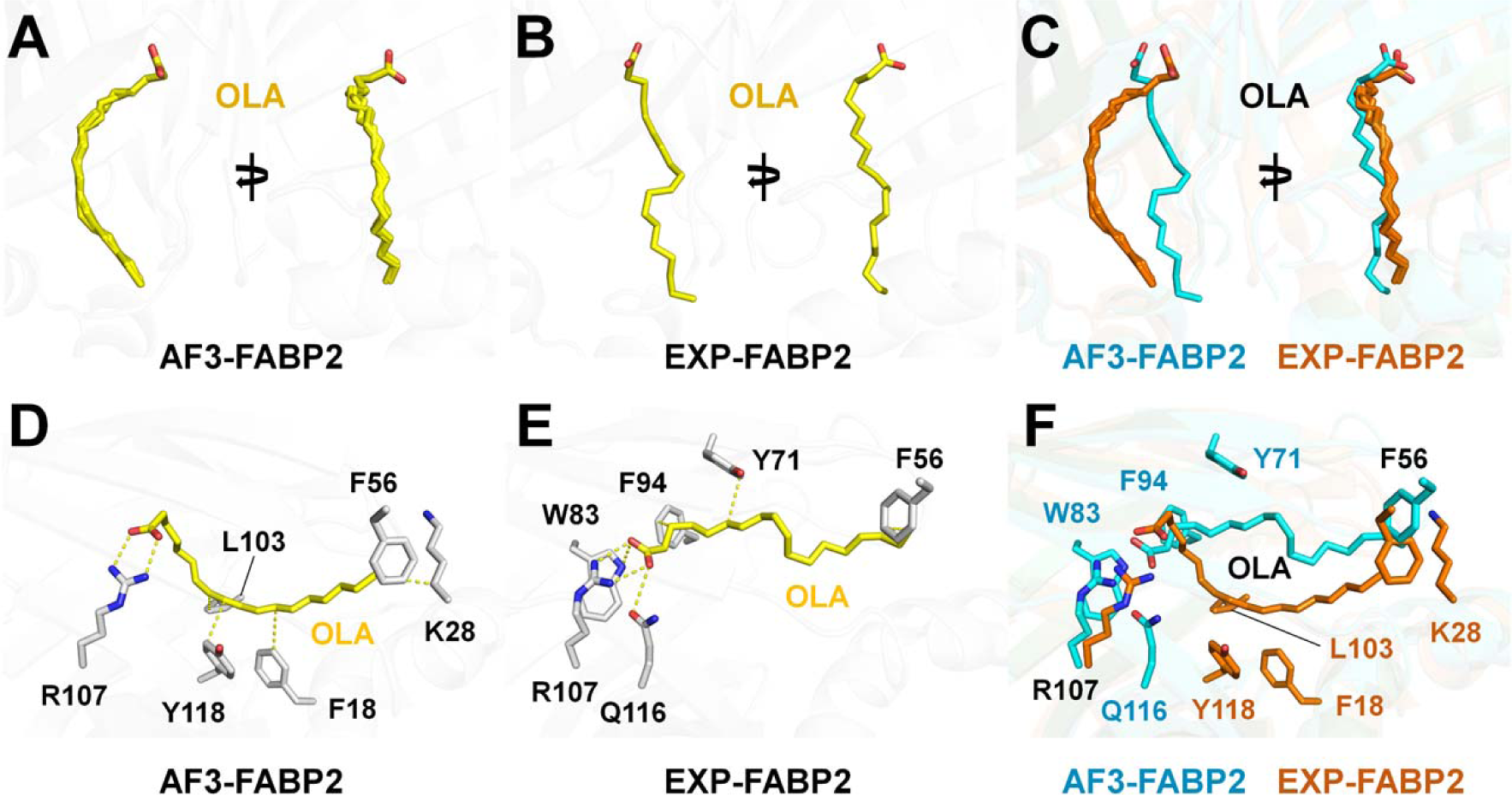
Structural comparison of OLA in AF3-FABP2 and EXP-FABP2. (A) Superimposition of five OLA ligands from AF3-FABP2. (B) OLA on EXP-FABP1 structure (PDB code: 2MO5). (C) Superimposition of OLA ligands from AF3-FABP2 (orange) and EXP-FABP2 (cyan) structures. Interactions of OLA ligands with (D) AF3-FABP2 and (E) EXP-FABP2 (PDB code: 2MO5) structures. (F) Superimposition of the OLA-binding site of AF3-FABP2 and EXP-FABP2 structures.

#### 3.2.3. FABP3

FABP3 (heart-type FABP or H-FABP) is abundantly expressed in skeletal and heart muscles [32]. FABP3 regulates the solubility, mobility, and utilization of FAs and muscle aging [33]. In PDB, the crystal structures of FABP3 complexed with MYR (4TKH and 7FZQ), PLM (2HMB, 4TKJ, 5B27, 5B28, 5B29, 6AQ1, 7FFK, and 7V2G), and OLA (5CE4, 1HMS, 7WE5, and 8GEW) have been deposited. These FA ligands were docked into FABP3 using AF3 and compared with experimental structures.

In MYR-docked AF3-FABP3, all MYR exhibited a U-shaped conformation, the carboxyl group and tail of the aliphatic chain were oriented toward the inside of the FA-binding pocket, and the middle of the aliphatic chain was oriented toward the FA-binding entrance. The atoms from the carboxyl group to the C6 atom in the aliphatic chain of MYR are positioned nearly identically, whereas the other atoms in the aliphatic chain showed slightly different positions (Figure 6A). In MYR-bound EXP-FABP3, two MYR showed similar U-shaped conformations, but the atom from C9 to the tail the aliphatic chain of MYR showed the instinct position (Figure 6B). Superimposition of MYR ligands from AF3-FABP3 and EXP-FABP3 structures showed a distinct position for the aliphatic chain of MYR, with an RMSD of 0.192–0.603 Å (Figure 6C). In the MYR-docked AF3-FABP3 structure, the carboxyl group of MYR interacted with Arg127 (2.73 Å), and the aliphatic chain of MYR was stabilized by hydrophobic interactions with Ala34 (3.64 Å), Pro39 (3.77 Å), and Lys59 (3.91 Å) (Figure 6D). In the MYR-bound EXP-FABP3 structure (PDB code: 4TKH), the carboxyl group of MYR interacted with Arg127 (2.75 and 3.08 Å), and the aliphatic chain of MYR was stabilized by hydrophobic interactions with Ala34 (3.97 Å) (Figure 6E). Superimposition of the MYR-binding site of AF3-FABP3 and EXP-FABP3 exhibited identical position and interaction of MYR with FABP3 (Figure 6F).

**Figure 6.**
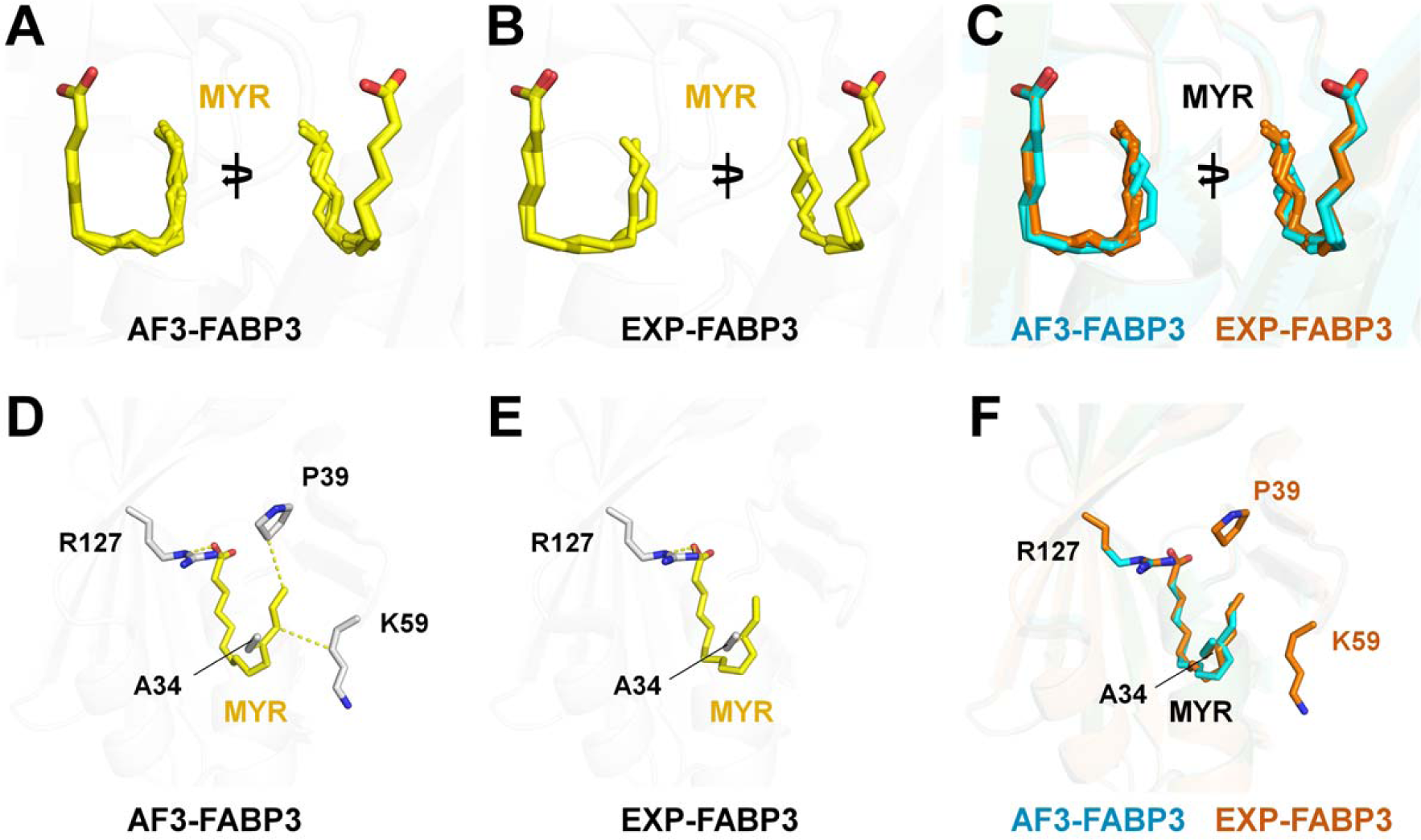
Structural comparison of MYR in AF3-FABP3 and EXP-FABP3 structures. (A) Superimposition of five MYR ligands from the AF3-FABP3 structure. (B) MYR on the EXP-FABP3 structure (PDB code: 4TKH and 7FZQ). (C) Superimposition of MYR ligands from AF3-FABP3 (orange) and EXP-FABP3 (cyan) structures. Interactions of MYR with (D) AF3-FABP3 and (E) EXP-FABP3 (PDB code: 4TKH) structures. (F) Superimposition of the MYR-binding site in AF3-FABP3 and EXP-FABP3.

In PLM-docked AF3-FABP3 and PLM-bound EXP-FABP3, all PLM ligands exhibited a U-shaped conformation and had similar positions and binding configurations as those of MYR binding (Figure 7A and 7B). Superimposition of PLM ligands from AF3-FABP3 and EXP-FABP3 structures had an RMSD of 0.123–0.209 Å (Figure 7C). In PLM-docked AF3-FABP3, the carboxyl group of PLM interacted with Arg127 (2.72 and 3.20 Å), and the aliphatic chain of PLM was stabilized by hydrophobic interactions with Tyr20 (3.89 Å), Leu24 (3.96 Å), Thr54 (3.41 Å), Phe58 (3.91 Å), Lys59 (3.95 Å), Asp77 (3.89 Å), and Leu116 (3.95 Å) (Figure 7D). In PLM-bound EXP-FABP3 (PDB code: 7FFK), the carboxyl group of PLM interacted with Arg126 (2.76 and 3.03Å) and Tyr128 (2.64 Å) and was stabilized by water bridges with Thr40 (2.69 Å) and Arg106 (2.88 Å). The aliphatic chain of PLM was stabilized by hydrophobic interactions with Phe16 (3.09 Å), Thr29 (3.96 Å), Ala33 (3.89 Å), Pro38 (3.99 Å), Phe57 (3.93 Å), Lys58 (3.89 Å), Ala75 (3.99 Å), and Asp76 (4.00 Å) (Figure 7E). Superimposition of the PLM-binding site of AF3-FABP3 and EXP-FABP3 exhibited that the position of PLM was mostly similar, and the PLM in EXP-FABP3 had a more extended interaction with FABP3 than the PLM in AF3-FABP3 (Figure 7F).

**Figure 7.**
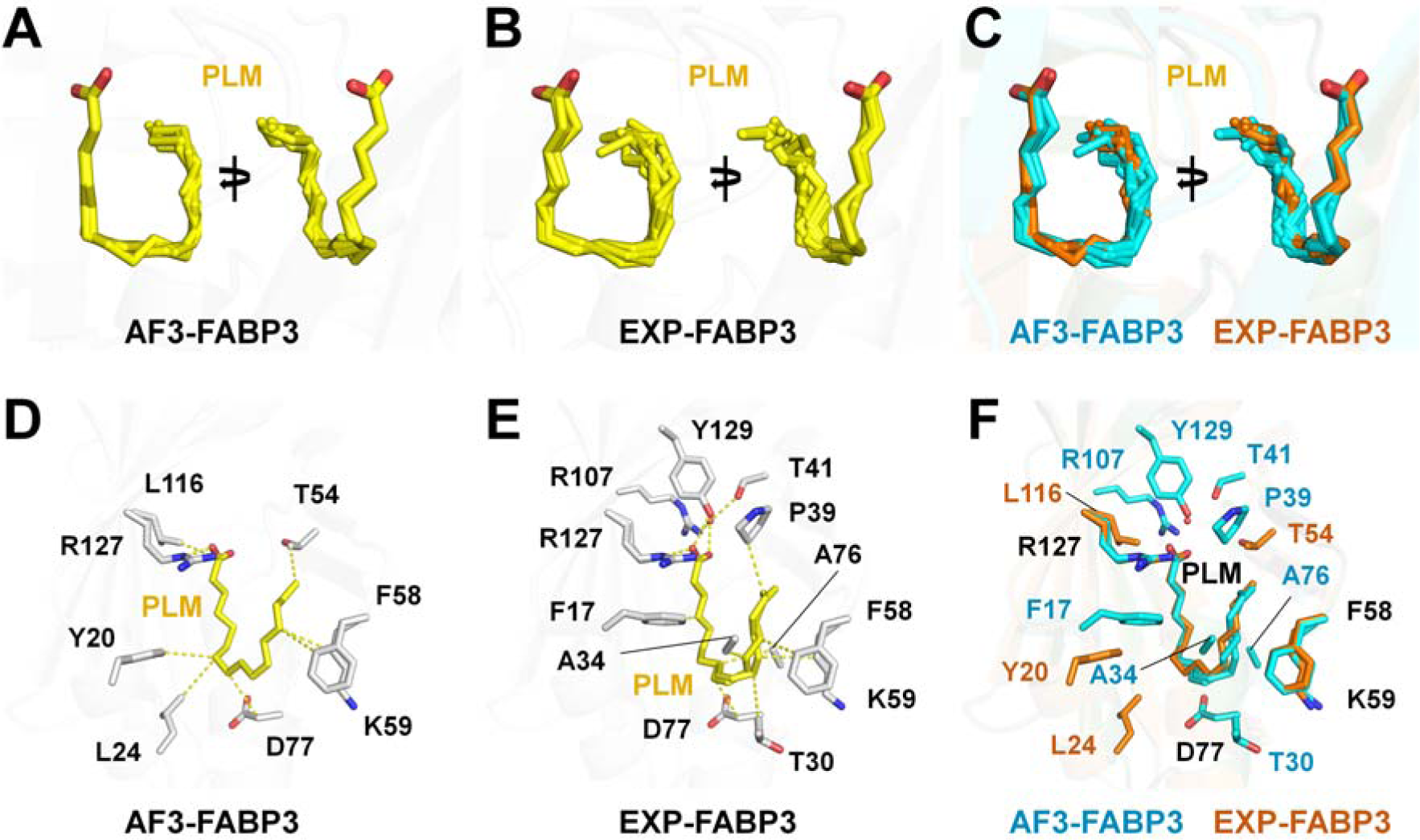
Structural comparison of PLM in AF3-FABP3 and EXP-FABP3 structures. (A) Superimposition of five PLM ligands from AF3-FABP3. (B) PLM on the EXP-FABP3 structure (PDB codes: 2HMB, 4TKJ, 5B27, 5B28, 5B29, 6AQ1, 7FFK, and 7V2G). (C) Superimposition of PLM ligands from the AF3-FABP3 (orange) and EXP-FABP3 (cyan) structures. Interactions of PLM with (D) AF3-FABP3 and (E) EXP-FABP3 (PDB code: 7FFK) structures. (F) Superimposition of the PLM-binding site of AF3-FABP3 and EXP-FABP3.

In OLA-docked AF3-FABP3 and OLA-bound EXP-FABP3, all PLM exhibited a U-shaped conformation, with similar positions and binding configurations as those of PLM binding (Figure 8A and 8B). Superimposition of PLM ligands from AF3-FABP3 and EXP-FABP3 structures had an RMSD of 0.132–0.680 Å (Figure 8C). In OLA-docked AF3-FABP3, the carboxyl group of PLM interacted with Arg127 (2.72 and 3.16 Å) and Tyr129 (2.48 Å). The aliphatic chain of OLA was stabilized by hydrophobic interactions with Tyr20 (3.80 Å), Leu24 (3.89 Å), Thr54 (3.61 Å), Phe58 (3.86 Å), Lys59 (3.93 Å), Thr61 (3.97 Å), and Asp77 (3.97 Å) (Figure 8D). In OLA-bound EXP-FABP3 (PDB code: 7WE5), the carboxyl group of OLA interacted with Arg127 (2.81 and 2.98 Å) and was stabilized by water bridges with Arg107 (2.69 Å). The aliphatic chain of OLA was stabilized by hydrophobic interactions with Phe17 (3.97 Å), Val26 (3.92 Å), Ala34 (3.96 Å), Thr54 (3.77 Å), and Asp77 (3.85 Å) (Figure 8E). Superimposition of the OLA-binding site of AF3-FABP3 and EXP-FABP3 exhibited that the position of OLA and its interaction with FABP3 were mostly similar (Figure 8F).

**Figure 8.**
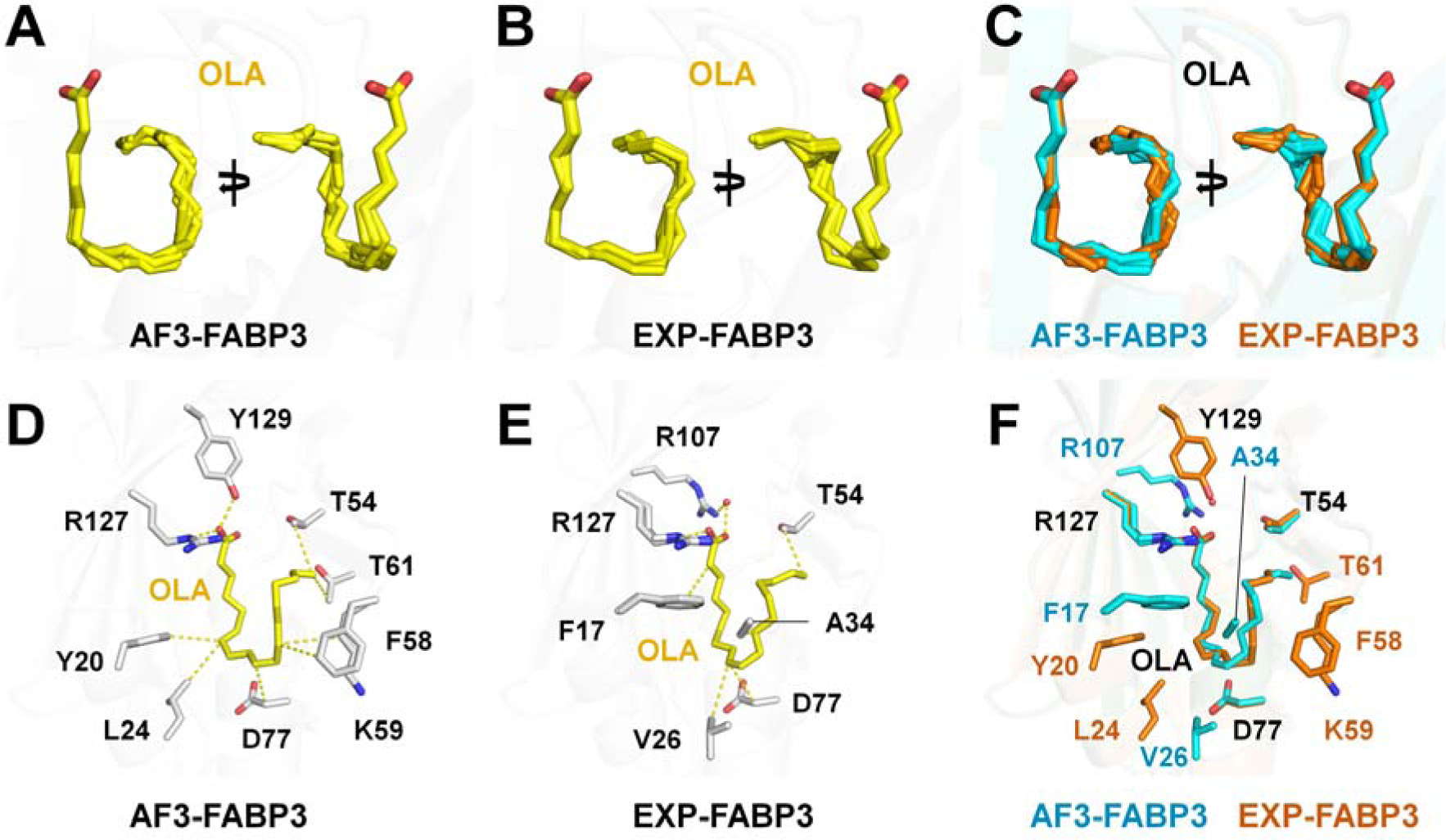
Structural comparison of OLA in AF3-FABP3 and EXP-FABP3 structures. (A) Superimposition of five OLA ligands from AF3-FABP3. (B) OLA on the EXP-FABP3 structure (PDB codes: 5CE4, 1HMS, 7WE5, and 8GEW). (C) Superimposition of OLA ligands from AF3-FABP3 (orange) and EXP-FABP3 (cyan) structures. Interactions of OLA with (D) AF3-FABP3 and (E) EXP-FABP3 (PDB code: 7WE5) structures. (F) Superimposition of the OLA-binding site of AF3-FABP3 and EXP-FABP3.

#### 3.2.4. FABP4

FABP4 (A-FABP/FABP4/aP2) is highly expressed in adipocytes and constitutes approximately 1% of all soluble proteins in adipose tissue [9]. FABP4 promotes lipid storage, distribution, transportation, decomposition, and metabolism, acting at the interface of metabolic and inflammatory pathways in adipocytes and macrophages [9, 34]. In PDB, the crystal structure of FABP4 complexed with PLM (2HNX) has been deposited. This was compared with PLM docking to FABP4 using AF3.

In PLM-docked AF3-FABP4, all PLM molecules were docked into almost identical positions with a U-shaped conformation in the FA-binding pocket of FABP4 (Figure 9A). The atoms from the carboxyl group of FA to C6 atom in the aliphatic chain of PLM were positioned nearly identically, whereas those from C6 atom to the tail of the aliphatic chain of FAs were at a slightly different position. In PLM-bound EXP-FABP4, PLM exhibited a U-shaped conformation in the FA-binding pocket of FABP4 (Figure 9B). The atomic positions from the carboxyl group to C4 in the aliphatic chain of PLM in EXP-FABP4 are almost identical to those in AF3-FABP4, but the other atomic region of PLM showed different positions between EXP-FABP4 and AF3-FABP4 (Figure 9C). In particular, the direction of the tail region (C14-C16) of the aliphatic chain of PLM is opposite between EXP-FABP4 and AF3-FABP4. Superimposition of PLM ligands from AF3-FABP4 and EXP-FABP4 structure had an RMSD of 0.353–0.690 Å. In PLM-docked AF3-FABP4, the carboxyl group of PLM interacted with Arg127 (2.72 and 3.33 Å) and Tyr129 (2.55 Å). The aliphatic chain of PLM was stabilized by hydrophobic interactions with Phe17 (3.51 Å), Tyr20 (3.72 Å), Ala34 (3.88 Å), Pro39 (3.78 Å), Ala76 (3.61 Å), and Ile105 (3.88 Å) (Figure 9D). In PLM-bound EXP-FABP4 (PDB code: 2HNX), the carboxyl group of PLM interacted with Arg127 (2.73 and 3.32 Å) and was stabilized by water bridges with Arg107 (2.84 Å) and Tyr129 (2.89 Å). The aliphatic chain of PLM was stabilized by hydrophobic interactions with Phe17 (3.66 Å), Ala34 (3.75 Å), Pro39 (3.64 Å), Phe58 (3.98 Å), and Asp77 (3.54 Å) (Figure 9E). Superimposition of the PLM-binding site of AF3-FABP4 and EXP-FABP4 exhibited that the position of PLM and its interaction with FABP3 were similar (Figure 9F).

**Figure 9.**
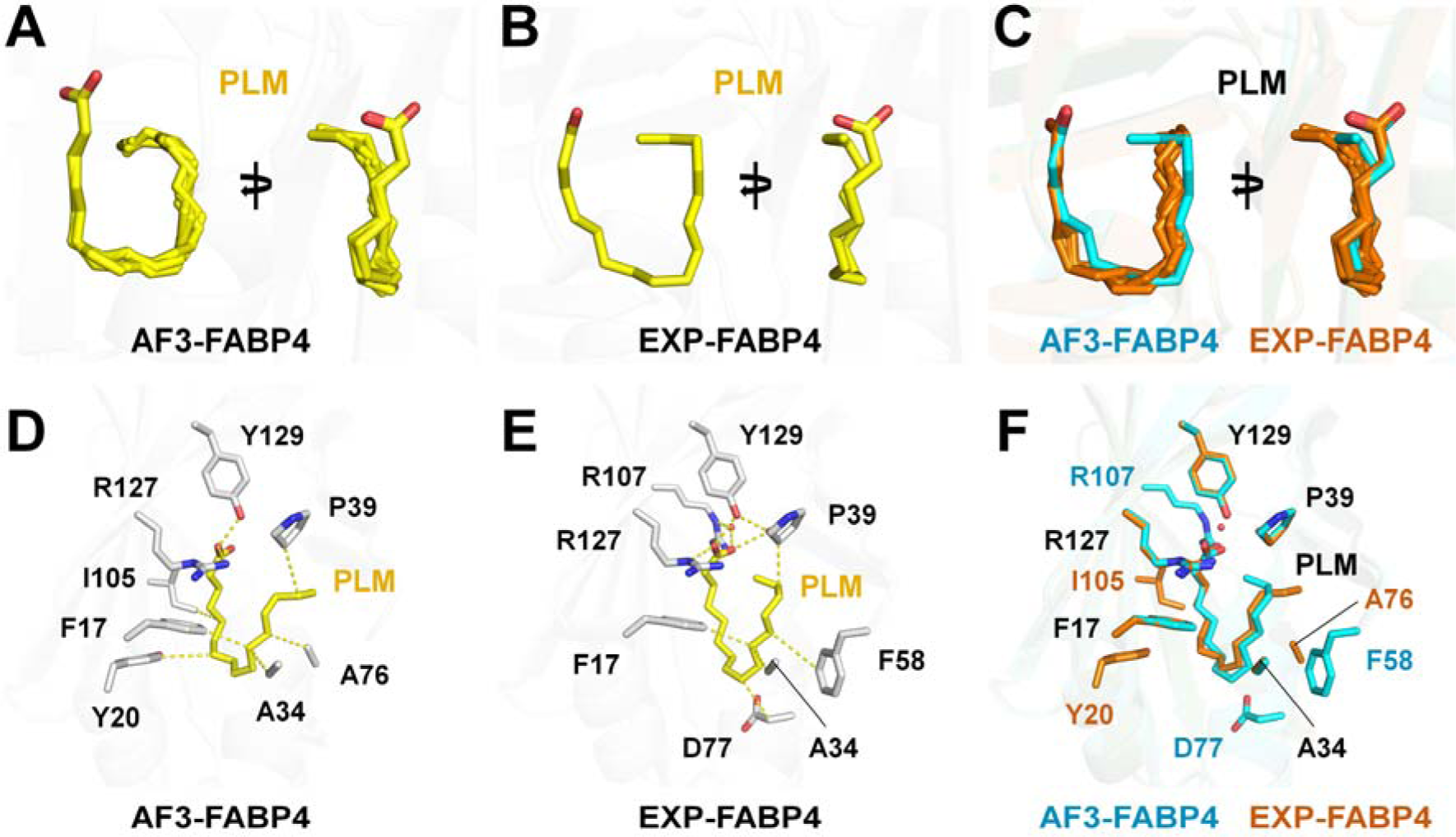
Structural comparison of PLM in AF3-FABP4 and EXP-FABP4. (A) Superimposition of five PLM ligands from AF3-FABP4. (B) PLM on the EXP-FABP4 structure (PDB code: 2HNX). (C) Superimposition of PLM ligands from AF3-FABP4 (orange) and EXP-FABP4 (cyan) structures. Interactions of PLM with (D) AF3-FABP and (E) EXP-FABP4 (PDB code: 2HNX) structures. (F) Superimposition of the PLM-binding site of AF3-FABP4 and EXP-FABP4.

#### 3.2.5. FABP5

FABP5 (epidermal FABP or E-FABP) is expressed in the skin, adipose tissue, and macrophage and participates primarily in FA uptake, transport, and metabolism in the cytoplasm [35]. In PDB, the crystal structure of FABP5 complexed with PLM (PDB code: 1B56) has been deposited. This was compared with PLM docking to FABP5 using AF3.

In PLM-docked AF3-FABP5, all PLM ligands docked into almost identical positions in the FA-binding pocket of FABP4 with a U-shaped conformation (Figure 10A). The atoms from the carboxyl group of FA to C7 atom in the aliphatic chain of PLM positioned nearly identically, whereas from C11 atom to the tail of the aliphatic chain of FAs showed large differences. In PLM-bound EXP-FABP5, the PLM ligand exhibited a C-shaped conformation in the FA-binding pocket of FABP5 (Figure 10B). The atomic positions from the carboxyl group of PLM in EXP-FABP5 were similar to those in AF3-FABP5, but the other atomic position of PLM in EXP-FABP5 showed differences with those in AF3-FABP5 (Figure 10C). Superimposition of PLM ligands from AF3-FABP5 and EXP-FABP5 had an RMSD of 0.866–1.488 Å. In PLM-docked AF3-FABP5, the carboxyl group of PLM interacted with Arg109 (3.87 Å), Arg129 (2.65 and 3.39 Å), and Tyr131 (2.51 Å). The aliphatic chain of PLM was stabilized by hydrophobic interactions with Phe19 (3.91 Å), Pro41 (3.95 Å), Thr56 (3.68 Å), and Asp79 (3.85 Å) (Figure 10D). In PLM-bound EXP-FABP5 (PDB code: 1B56), the carboxyl group of PLM interacted with Arg109 (3.43 Å), Arg129 (3.42 Å), and Tyr131 (2.81 Å) and was stabilized by water bridges with Arg109 (2.68 Å). The aliphatic chain of PLM was stabilized by hydrophobic interactions with Ala78 (3.86 Å) and Val118 (3.29 Å) (Figure 10E). Superimposition of the PLM-binding site of AF3-FABP5 and EXP-FABP5 exhibited significantly different positions of the aliphatic chain of PLM and its interacting residues (Figure 10F).

**Figure 10.**
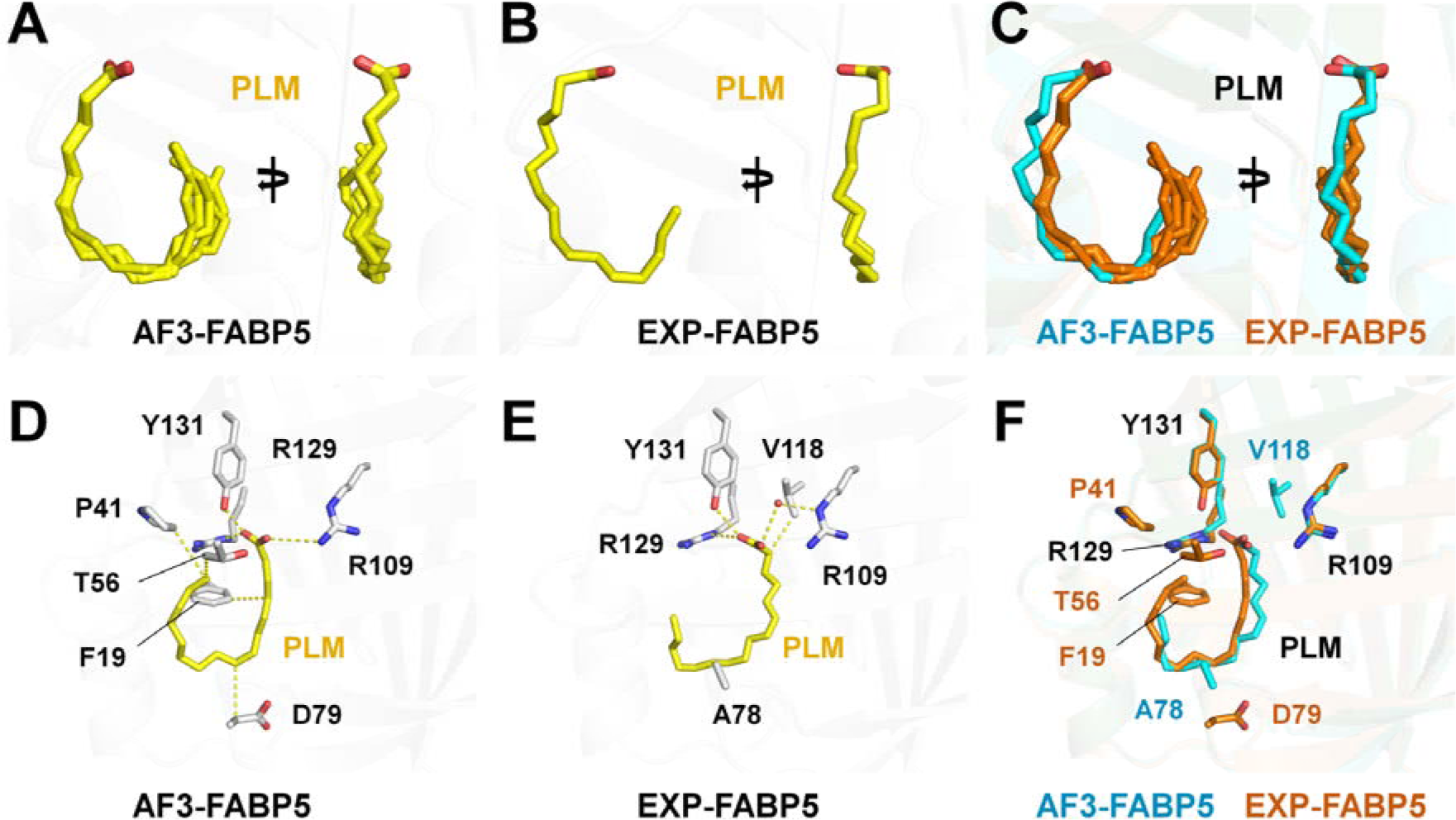
Structural comparison of PLM in AF3-FABP5 and EXP-FABP5. (A) Superimposition of five PLM ligands from AF3-FABP5. (B) PLM on the EXP-FABP5 structure (PDB code: 1B56). (C) Superimposition of PLM ligands from AF3-FABP5 (orange) and EXP-FABP5 (cyan) structures. Interactions of PLM with (D) AF3-F€5 and (E) EXP-FABP5 (PDB code: 1B56) structures. (F) Superimposition of the PLM-binding site of AF3-FABP5 and EXP-FABP5.

#### 3.2.6. FABP7

FABP7 (B-FABP) is abundant in the embryonic brain and is expressed in astrocytes and neural progenitors [3]. FABP7 is involved in various cellular functions, such as the uptake and transport of lipids and protein metabolism [36, 37]. FABP7 has high affinity to docosahexaenoic acid and a polyunsaturated FA, which are both abundant in the brain [38, 39]. FABP7 is important for neurodevelopment and the central nervous system [40]. In PDB, the crystal structure of PLM-bound FABP7 (PDB code: 7E25) and OLA-bound FABP7 (PDB code: 1FE3) have been deposited. These were compared with the structure PLM or OLA docked to FABP7 using AF3.

In PLM-docked AF3-FABP7, all PLM ligands docked into the FA-binding pocket of FABP7 almost identical positions with a U-shaped conformation (Figure 11A). The atoms from the carboxyl group of FA to C7 atom in the aliphatic chain of PLM positioned nearly identically, whereas from CB atom to the tail of the aliphatic chain of FAs showed large differences. In PLM-bound EXP-FABP7, the PLM ligand exhibited a U-shaped conformation in the FA-binding pocket of FABP7 (Figure 11B), but the PLM binding configuration was significantly different with PLM binding in AF3-FABP7. The superimposition structure showed that atomic positions from the carboxyl group to the aliphatic tail of PLM in EXP-FABP7 were almost different to those in AF3-FABP7 (Figure 11C). Superimposition of PLM ligands on AF3-FABP7 and EXP-FABP7 had similarity with an RMSD of 0.938–0.970 Å. In PLM-docked AF3-FABP7, the carboxyl group of PLM interacted with Arg127 (2.81 and 3.22 Å) and Tyr129 (2.43 Å), and the aliphatic chain of PLM was stabilized by hydrophobic interactions with Pro39 (3.82 Å), Phe58 (3.83 Å), and Asp77 (4.00 Å) (Figure 11D). In PLM-bound EXP-FABP (7E25), the carboxyl group of PLM interacted with Tyr129 (2.51 Å) and Arg127 (2.82 and 3.31 Å), and the aliphatic chain of PLM was stabilized by hydrophobic interactions with Leu24 (3.98 Å), Thr30 (3.85 Å), Thr37 (3.81 Å), Pro39 (3.93 Å), Thr54 (3.98 Å), Phe58 (3.83 Å), and Asp77 (3.77 Å) (Figure 11E). Superimposition of the PLM-binding site of AF3-FABP7 and EXP-FABP7 exhibited that the position of PLM was different, and the PLM in EXP-FABP7 had a more extended interaction with FABP7 than the PLM in AF3-FABP7 (Figure 11F).

**Figure 11.**
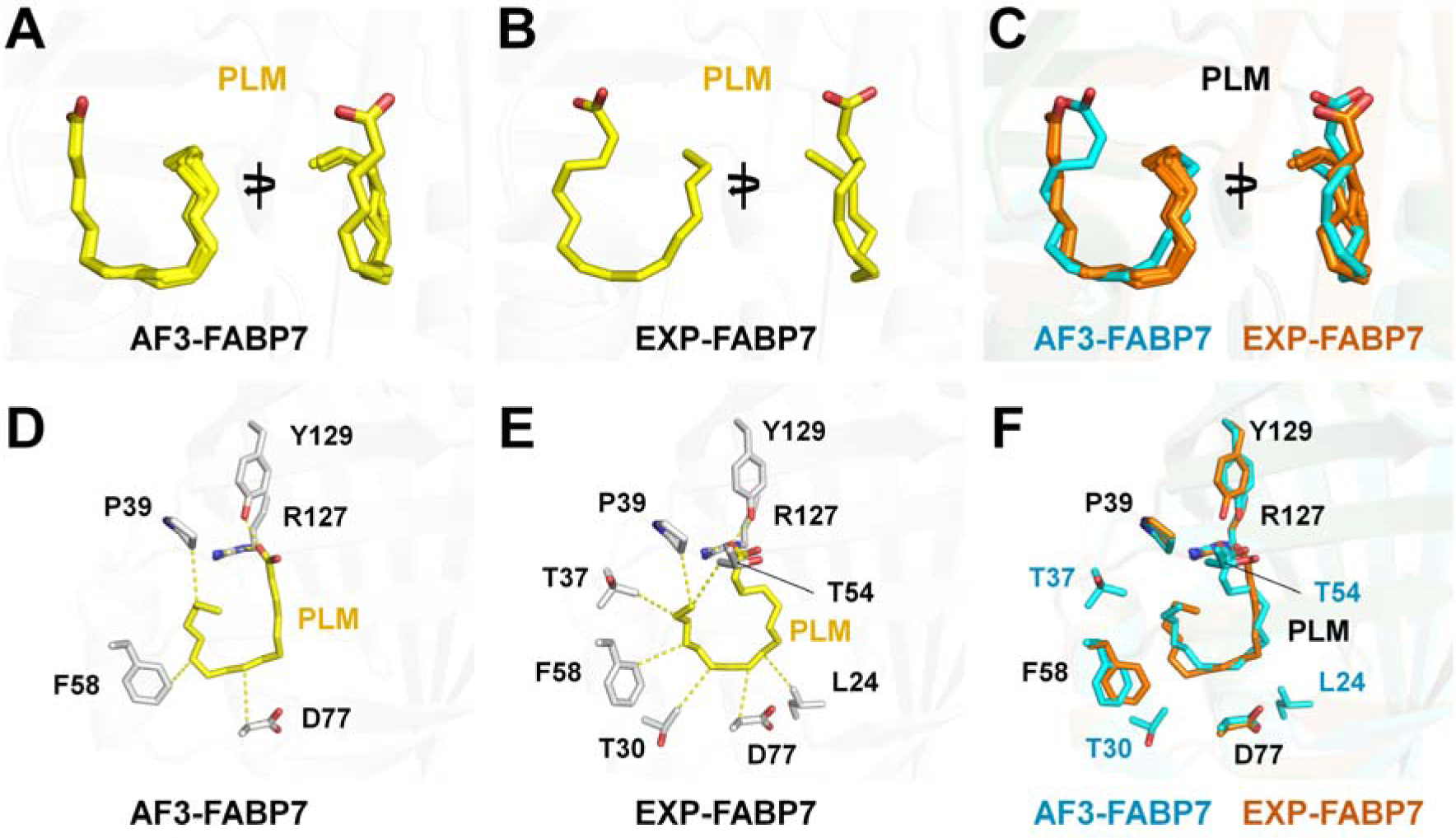
Structural comparison of PLM in AF3-FABP7 and EXP-FABP7. (A) Superimposition of five PLM ligands from AF3-FABP7. (B) PLM in the EXP-FABP7 structure (PDB code: 7E25). (C) Superimposition of PLM ligands from AF3-FABP7 (orange) and EXP-FABP7 (cyan) structures. Interactions of PLM with (D) AF3-FABP7 and (E) EXP-FABP7 (PDB code: 7E25) structures. (F) Superimposition of the PLM-binding site of AF3-FABP7 and EXP-FABP7.

In OLA-docked AF3-FABP7, all OLA ligands docked into almost similar positions showing mainly U-shaped conformation. The aliphatic tail of OLA ligands oriented toward the hydrophobic region in the FA-binding pocket of FABP7 (Figure 12A). The atoms from the carboxyl group of FA to C5 in the aliphatic chain of OLA positioned nearly identically, whereas the tail (C14–C18) of the aliphatic chain of FAs showed a different conformation. In OLA-bound EXP-FABP7, the OLA ligand mainly exhibited a U-shaped conformation in the FA-binding pocket of FABP7 (Figure 12B), but the carboxyl group and tail of OLA showed distinct conformations from those in AF3-FABP7 (Figure 12C). The orientations of the OLA ligand with a U-shaped conformation in the FA-binding pocket from AF3-FABP7 and EXP-FABP7 differed. Superimposition of OLA ligands on AF3-FABP7 and EXP-FABP7 showed an RMSD of 0.529–0.690 Å.

**Figure 12.**
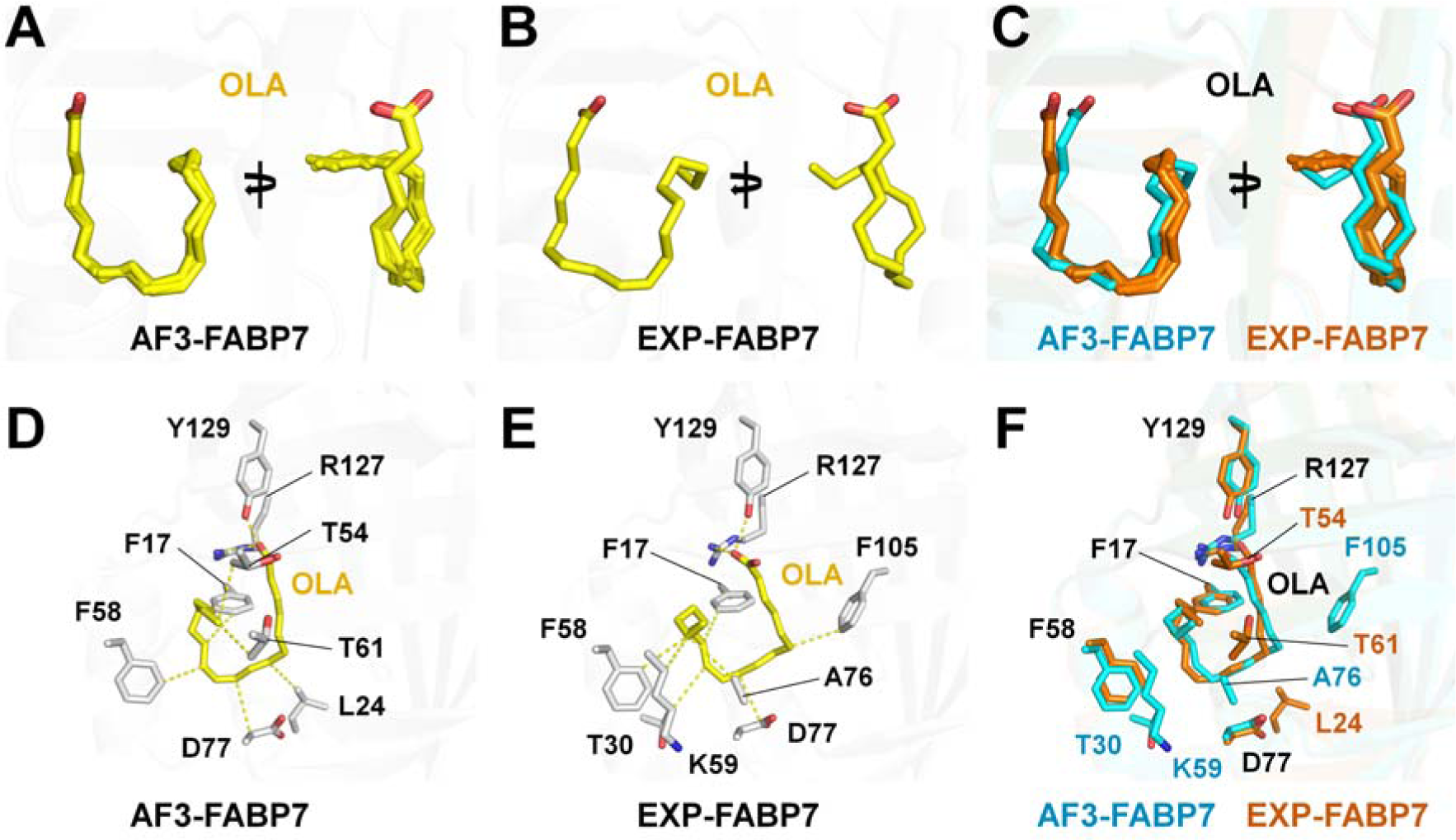
Structural comparison of OLA in AF3-FABP7 and EXP-FABP7. (A) Superimposition of five OLA ligands from AF3-FABP7. (B) OLA on the EXP-FABP7 structure (PDB code: 1FE3). (C) Superimposition of OLA ligands from AF3-FABP7 (orange) and EXP-FABP7 (cyan) structures. Interactions of OLA with (D) AF3-FABP7 and (E) EXP-FABP7 (PDB code: 1FE3) structures. (F) Superimposition of the OLA-binding site of AF3-FABP7 and EXP-FABP7.

In OLA-docked AF3-FABP7, the carboxyl group of OLA interacted with Arg127 (2.84 and 3.22 Å) and Tyr129 (2.33 Å), and the aliphatic chain of PLM was stabilized by hydrophobic interactions with Phe17 (3.98 Å), Leu24 (3.94 Å), Thr54 (3.76 Å), Phe58 (3.62 Å), Thr61 (3.73 Å), and Asp77 (3.98 Å) (Figure 12D). In OLA-bound EXP-FABP (PDB code: 1FE3), the carboxyl group of OLA interacted with Tyr129 (2.92 Å) and Arg127 (3.15 Å), and the aliphatic chain of OLA was stabilized by hydrophobic interactions with Phe17 (3.85 Å), Thr30 (3.86 Å), Phe58 (3.86 Å), Lys59 (3.81 Å), Ala76 (3.95 Å), Asp77 (3.60 Å), and Phe105 (3.76 Å) (Figure 12E). Superimposition of the OLA-binding site of AF3-FABP7 and EXP-FABP7 exhibited that the position and conformation of OLA were different (Figure 12F).

#### 3.2.7. FABP8

FABP8 (M-FABP, peripheral myelin protein 2, PMP2, myelin P2 protein) is expressed in vertebrate peripheral nervous system myelin and Schwann cells [41, 42]. FABP8 acts as a lipid carrier involved in the production and maintenance of myelin [43]. In PDB, the 29 crystal structures of PLM-bound FABP8 have been deposited, which were compared with the PLM-docked AF3-FABP8 structure.

In PLM-docked AF3-FABP8, all PLM ligands docked into almost identical positions in the FA-binding pocket of FABP8 with a U-shaped conformation (Figure 13A). The atoms from the carboxyl group of FA to C6 atom in the aliphatic chain of PLM positioned nearly identically, whereas those from C7 atom to the tail of the aliphatic chain of FAs showed differences. In PLM-bound EXP-FABP8, 29 PLM ligands with a U-shaped configuration showed various positions and diverse conformations of the aliphatic chain (Figure 13B). Superimposition of PLM ligands from AF3-FABP8 and EXP-FABP8 showed that the position of PLM overlapped (Figure 13C). Superimposition of PLM ligands from AF3-FABP8 and EXP-FABP8 (6S2M) with highest resolution structure showed an RMSD of 0.378–0.818 Å. In PLM-docked AF3-FABP8, the carboxyl group of PLM interacted with Arg107 (3.77 Å), Tyr129 (2.54 Å), and Arg127 (2.64 Å). The aliphatic chain of PLM was stabilized by hydrophobic interactions with Tyr20 (3.93 Å), Pro39 (3.85 Å), Thr54 (3.46 Å), Phe58 (3.77 Å), and Ala76 (3.71 Å), and Asp77 (3.79 Å) (Figure 13D). In PLM-bound EXP-FABP8 (PDB code: 6S2M), the carboxyl group of PLM interacted with Arg107 (3.05 Å) and Arg127 (2.88 Å). The aliphatic chain of PLM was stabilized by hydrophobic interactions with Phe17 (4.00 Å), Leu24 (3.94 Å), Thr30 (3.72 Å), Pro39 (3.97 Å), and Ala76 (3.97 Å) (Figure 13E). Superimposition of the PLM-binding site of AF3-FABP8 and EXP-FABP8 exhibited that the position and conformation of OLA were significantly different (Figure 13F). However, considering the various binding conformations of PLM in EXP-FABP8s (Figure 13B), the binding configuration of PLM in AF3-FABP8 were reliable.

**Figure 13.**
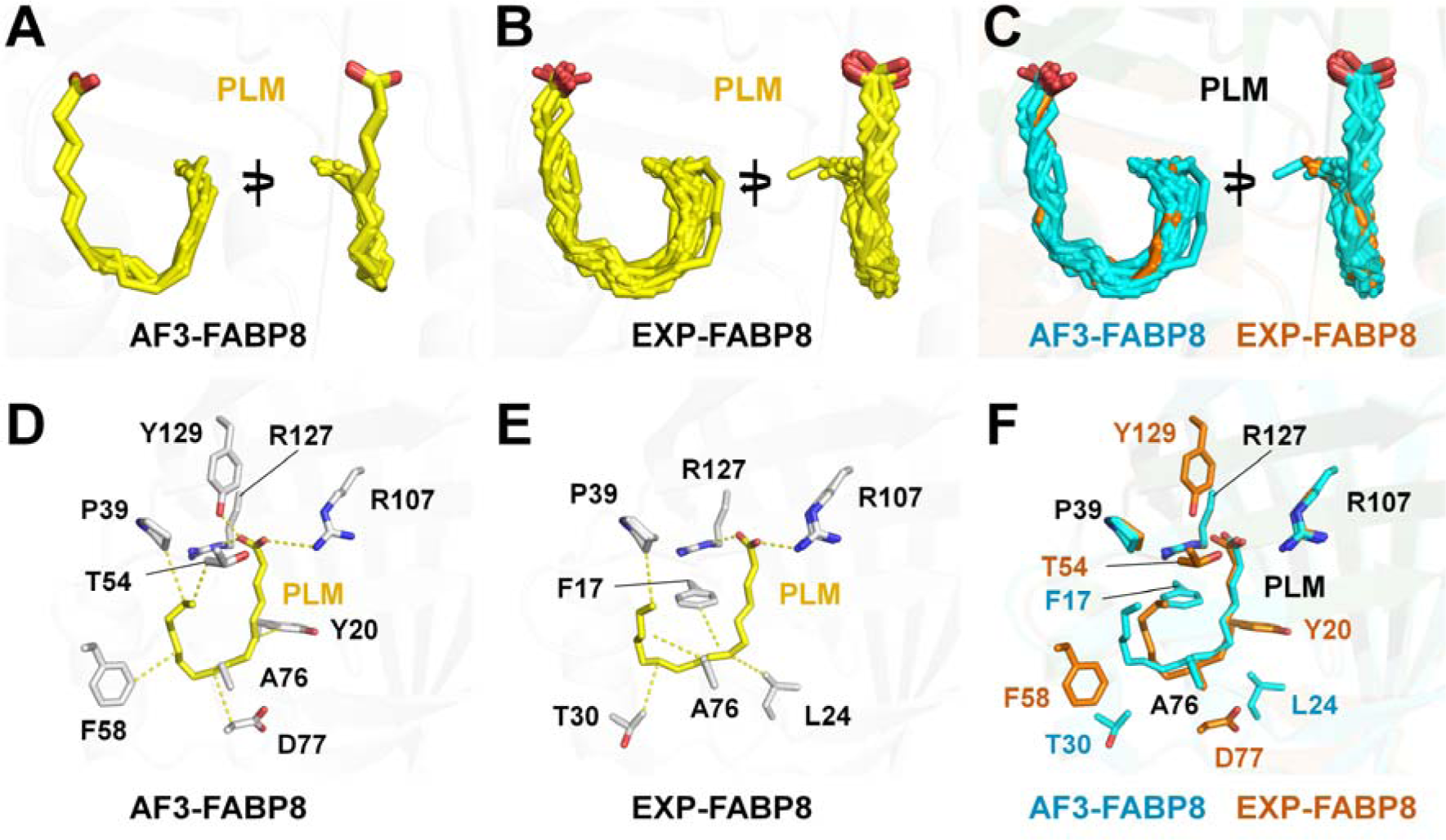
Structural comparison of PLM in AF3-FABP8 and EXP-FABP8. (A) Superimposition of the five PLM ligands from AF3-FABP8. (B) OLA in the EXP-FABP8 structure (PDB code: 6S2M). (C) Superimposition of PLM ligands from AF3-FABP8 (orange) and EXP-FABP8 (cyan) structures. Interactions of PLM with the (D) AF3-FABP8 and (E) EXP-FABP8 (PDB code: 6S2M) structures. (F) Superimposition of the PLM-binding site of AF3-FABP8 and EXP-FABP8.

#### 3.2.8. FABP9

FABP9 is expressed in the mammalian testis [44] and has a role in determining germ cell fate and preserving sperm quality [45] In PDB, the crystal structures of MYR-bound FABP9 have been deposited, which were compared with the MYR-docked AF3-FABP9 structure.

In MYR-docked AF3-FABP9, all MYR ligands docked into almost identical positions in the FA-binding pocket of FABP9 with a U-shaped conformation (Figure 14A). The atoms from the carboxyl group of FA to C7 atom in the aliphatic chain of MYR were positioned nearly identically, whereas those from C7 atom to the tail of the aliphatic chain of FAs showed differences. In MYR-bound EXP-FABP9, MYR was bound to the FA-binding site of FABP9 with an L-shaped configuration (Figure 14B). Superimposition of MYR from AF3-FABP8 and EXP-FABP8 showed that the positions from the carboxylate group to C7 atom of PLM were similar, but other aliphatic chain of MYR were significantly different (Figure 13C). Superimposition of MYR ligands from AF3-FABP9 and EXP-FABP9 (7FY1) showed an RMSD of 2.003–2.197 Å. In MYR-docked AF3-FABP9, the carboxyl group of MYR interacted with Arg127 (2.70 Å). The aliphatic chain of MYR was stabilized by hydrophobic interactions with Phe17 (3.67 Å), Ala34 (3.77 Å), Pro39 (3.84 Å), Phe58 (3.99 Å), and Ala76 (3.75 Å) (Figure 14D). In MYR-bound EXP-FABP9 (PDB code: 7FY1), the carboxyl group of MYR interacted with Tyr129 (2.61 Å), Arg107 (3.25 Å), and Arg127 (3.03 Å). The aliphatic chain of MYR was stabilized by hydrophobic interactions with Phe17 (3.98 Å), Leu24 (3.86 Å), Ala34 (3.52 Å), Gln59 (3.61 Å), Ala76 (3.93 Å), Asp77 (3.81 Å), and Ile105 (3.98 Å) (Figure 14E). Superimposition of the MYR-binding site of AF3-FABP9 and EXP-FABP9 showed significantly different MYR positions and conformations as well as MYR-interacting residues (Figure 14F).

**Figure 14.**
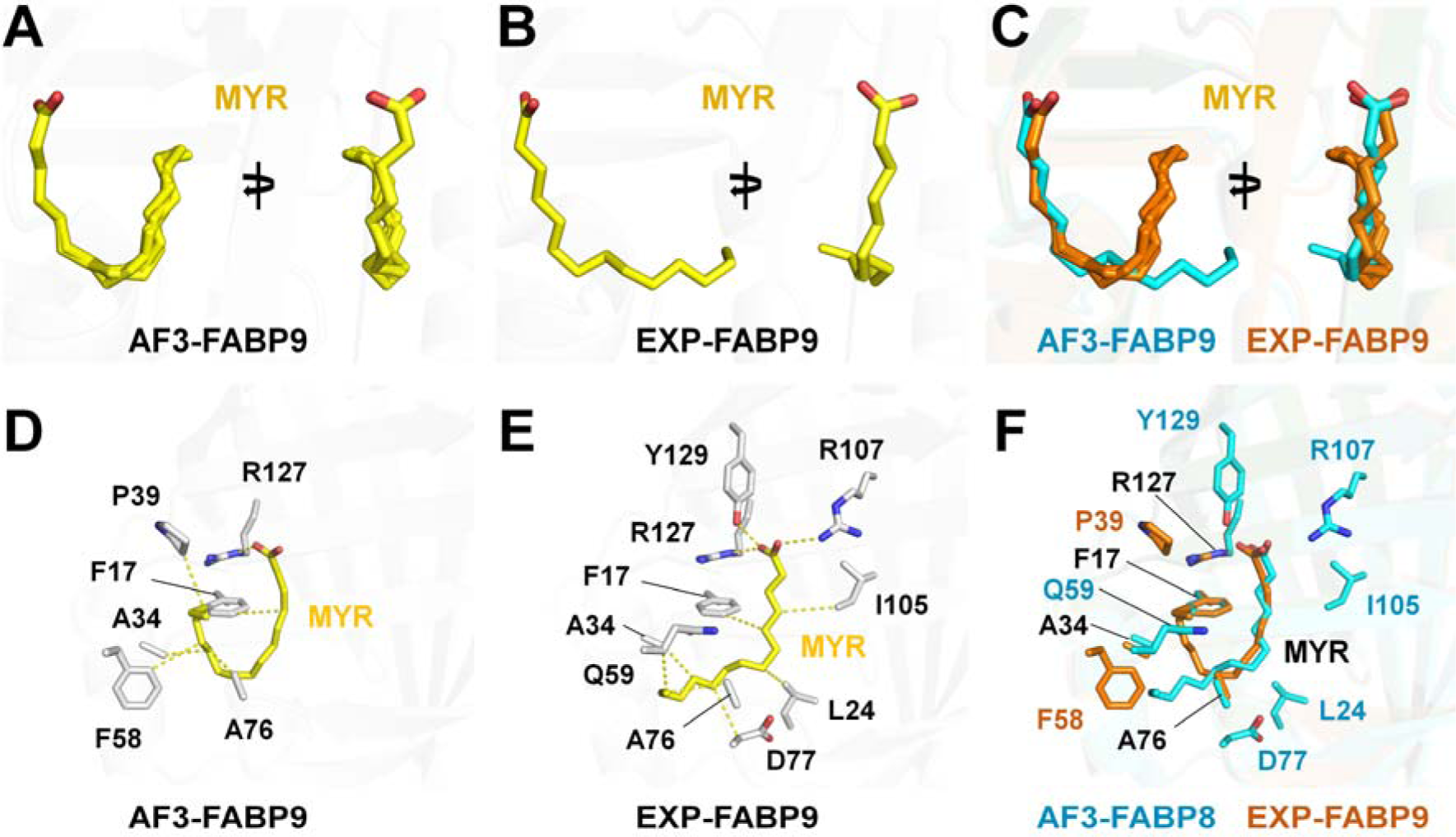
Structural comparison of MYR in AF3-FABP9 and EXP-FABP9. (A) Superimposition of five MYR ligands from AF3-FABP9. (B) MYR on EXP-FABP9 structure (PDB code: 7FY1). (C) Superimposition of MYR ligands from AF3-FABP9 (orange) and EXP-FABP9 (cyan) structure. Interactions of MYR with (D) AF3-FABP9 and (E) EXP-FABP9 (PDB code: 7FY1) structure. (F) Superimposition of the MYR-binding site of AF3-FABP9 and EXP-FABP9.

## 4. Discussion

Ligand docking studies play a critical role in understanding experimentally unproven protein– ligand interactions. They prove to be efficient and cost saving during inhibitor development and drug discovery. AF3 is a recently developed AI-based program that can predict interactions between proteins and ligands. Here, to evaluate the reliability of ligand docking results in AF3, FA-docked FABP structures generated using AF3 were compared with the experimental FA-bound FABP structures. The conformations and positions of FAs docked in FABP3–5 and FABP8 using AF3 showed similarity to the experimental structures. However, in FABP1, FABP2, FABP7, and FABP9, the structural characteristics of FAs docked into FABP using AF3 were significantly different from the experimental FA-bound FABP structures.

In the experimental two FA-bound FABP1, the binding conformation of U-shaped PLM and OLA at the first FA-binding site of FABP1 showed similarity to the two FA-docked AF3-FABP1 in terms of position and conformation. In the two FA-bound EXP-FABP1, the carboxyl group of PLM2 with a linear form at the second FA-binding site was oriented toward the inside of the FA-binding pocket. By contrast, the carboxyl group of OLA2 with a linear form at the second FA-binding site was oriented toward the entrance of the FA-binding pocket. The PLM and OLA docking results for FABP1 using AF3 showed that the orientation of the carboxyl group of some FAs was opposite to the experimental structures. This result indicates that verification of FA orientation may be necessary in FA docking models generated using AF3.

For FABP2, the position and configuration of the OLA docking model on FABP2 generated using AF3 were significantly different from the OLA in the experimental structure. OLA has a double bond between C8 and C9 atoms, preferring a *trans*-conformation at that position, which is consistent with the experimentally determined structure for OLA in FABP2. However, the double bond region of OLA docked into FABP2 exhibited a *cis*-conformation using AF3. Accordingly, when verifying the OLA docking results generated with AF3, it is necessary to check whether the inherently preferred *trans*-conformation of OLA is maintained.

For FABP7 and FABP9, the conformation and position of the FAs docked into FABP using AF3 were different from those of experimental FA-bound FABP. This indicated that the amino acids interacting with FA and their conformations varied and that FA conformation and position affected the structural information of FA-interacting amino acids. The position and conformation of the FA docking model in FABP3–5 and FABP8 were similar with the experimental structures of FA bound to FABPs. However, its interacting between FA and FABP were significantly different excluding the MYR-docked AF3-FABP3. In other words, even if the FA docking position or conformation for FABP provided using AF3 was similar to the experimental results, there was a difference in detailed interactions, indicating that the verification of the interacting amino acid is necessary. This may especially affect the results of follow-up studies, such as studies on the design of FABP inhibitors, based on docking results.

In FA docked to FABP using AF3, FA was docked into the native FABP structure, providing information solely about the position of FA and its interaction with the protein. By contrast, experimental FABP structures included details about FA interactions with amino acids within the FA-binding pocket as well as interactions with water molecules. For instance, in FABP1– PLM, FABP3–PLM, FABP3–OLA, FABP4–PLM, and FABP5–PLM, the carboxyl groups of FAs were stabilized by water bridges connected to the main chain or side chains of amino acids within the FA-binding pocket of FABP. This suggests that water molecules inside the FA-binding pocket of FABP may influence the conformation and position of FA. Because AF3 does not include information about these water molecules, the docking results may differ. By contrast, the experimental structure provides valuable information about the positions of water molecules, which can be crucial for inhibitor design and understanding FA recognition. Currently, AF3 structures do not incorporate details about the water molecules.

FABPs can bind not only FAs but also other molecules. For instance, in the MYR-bound FABP4 mutant (PDB codes: 7FYH and 7G0X), 6-fluoro-1,3-benzothiazol-2-amine or isoquinolin-3-amine bind to the FA-binding pocket. These molecules affect the conformation of FA when compared with MYR-bound FABP4 mutant alone. This indicates that when docking FA to FABP, the position or binding conformation of FA may change if another molecule is present in the FA-binding site. This consideration should apply not only to AF3 but also to other docking programs.

In experimentally determined FABP1–PLM, FABP3–PLM, FABP3–OLA, and FABP8–PLM, slightly different conformations and positions of FA ligands were observed across multiple experimental structures. These subtle changes in the position and conformation of the FAs may vary depending on protein crystallization conditions, data collection environment, quality of electron density maps, or natural properties of FAs binding to FABP. This finding indicates that the conformation of FAs within FABP may vary. If there are diverse experimental models of FAs, there may also be differences in the interpretation of FA docking results in AF3. Therefore, to accurately analyze FA docking using AF3, having a greater variety of FABP structures allows for more precise comparative analysis.

## 5. Conclusion

The results of docking FAs to FABP using AF3 were compared with the experimental structure. It structurally explained discrepancies and similarities between FA docking results and experimental structures. These findings provide insights into the understanding and future applications of ligand docking based on FA using AF3.

## Author Contributions

Ki Hyun Nam wrote the manuscript.

## Data availability

The code used to run the docking, measure the distances, and create the figures

